# The N-terminus of *Plasmodium falciparum* Circumsporozoite Protein Contains Three Non-Overlapping Murine B-cell Epitope Regions

**DOI:** 10.1101/2024.01.18.574923

**Authors:** Nathan Beutler, Elijah Garcia, Yen-Chung Lai, Justin Ndihokubwayo, Kiara Gambuzza, Jerry Zhao, Dennis R. Burton, Thomas F. Rogers

**Affiliations:** Department of Immunology and Microbiology, The Scripps Research Institute, La Jolla, CA 92037, USA; Mayo Clinic Medical Scientist Training Program, Mayo Clinic Alix School of Medicine, Rochester, MN; Division of Infectious Diseases, Department of Medicine, University of California, San Diego, La Jolla, CA 92037, USA; IAVI Neutralizing Antibody Center, The Scripps Research Institute, La Jolla, CA 92037, USA; Ragon Institute of Massachusetts General Hospital, Massachusetts Institute of Technology and Harvard University, Cambridge, MA 02139, USA

## Abstract

The generation of an anti-malarial vaccine that produces broad, potent, and durable responses is highly desirable to control the burden of *Plasmodium falciparum* disease. Current vaccines have offered modest efficacy ranging from 50%-70%, likely associated with antibody responses that are relatively short lived and strain specific. Currently approved malaria vaccines, RTS,S and R21, target the repeat region and C-terminal region of *Plasmodium falciparum* CSP, leaving the N-terminal region of CSP neglected as a target for protective immunogen design. Here, we isolate and express a panel of memory B-cell derived N-terminal CSP-specific monoclonal antibodies (mAbs) from mice immunized with an N-terminal CSP specific immunogen. The characterization of N-terminal specific mAbs including peptide walking and affinity experiments indicate that these antibodies target three distinct sites within the N-terminus of CSP. Site ntCSP-A contains the Region I (RI) cleavage site, which has been previously defined, whereas the remaining two sites are in previously undescribed locations upstream of RI, termed ntCSP-B and ntCSP-C.

## Introduction

Malaria remains a global threat, leading to 608,000 deaths and 249 million infections worldwide in 2022^1^. Malaria disproportionately affects children and pregnant woman, who make up the majority of malaria deaths^1^. Malaria disease is caused by the *Plasmodium* parasite, which is spread by a bite from an infected female anopheles mosquito. When the infected mosquito feeds, it releases sporozoites into the human host bloodstream, which later infect host hepatocytes and through different maturation processes, host red blood cells. The symptomatic stage of infection can include fever, chills, headache, fatigue, confusion, seizures, difficulty breathing, and death^1^. Many medical countermeasures have been developed to curb the burden of malaria including the widespread use of mosquito nets, anti-malarial drugs, monoclonal antibody therapy, and vaccines. Unfortunately, malaria continues to show increased resistance to anti-malarial drugs in WHO-defined African regions, which account for 95.4% of all malaria related deaths in 2022^1-3^. To slow the development of drug resistance and limit overall malaria disease and death, the development of an effective malarial vaccine has been prioritized^4^.

The leading anti-malarial vaccine, RTS,S/AS01, showed modest efficacy around 50%, which was short lived in phase 3 clinical trials in Africa^5,6^. Nevertheless, the vaccine was recommended for use in children under the name “Mosquirix”. RTS,S is composed of a virus-like particle containing 19 NANP repeats and the C-terminal region of the lab-adapted 3D7 strain of *Plasmodium falciparum* circumsporozoite surface protein (*pf*CSP) linked to hepatitis B surface antigen(HBsAg)^7^. RTS,S trial participants elicited antibodies toward both the NANP repeat region and C-terminal region of CSP, both of which were associated with protection^8-11^. The efficacy of the vaccine decreased over time, which was associated with waning antibody titers^5,6,12^. A similar leading vaccine candidate, R21, is composed of the same HBsAg-CSP but without unmodified HbsAg that is part of RTS,S vaccine and paired with a Matrix-M adjuvant^13^. In a follow up of a recent phase 2 clinical trial where R21 was administered to children in Burkina Faso, efficacies of 71% and 80% were reported in the low- and high-dose adjuvant groups respectively, 24 months after dose 3^14^. In a similar fashion to RTS,S, antibody titers also quickly waned over time to almost baseline levels prior to the 4th dose^14,15^.

*pfCSP* is composed of an N-terminal region, a junctional region, a central NANP repeat region, and a C-terminal region^16^. The C-terminal region is composed of a α-thrombospondin repeat domain that has been suggested to be involved in sporozoite binding to heparan sulfate on hepatocytes and invasion ^17-19^. This region has been shown to contain two protective B-cell targets, α-ctCSP, which targets non-conserved regions within Th2R and Th3R and β-ctCSP, which targets the highly conserved RII+ and CS.T3^20^ regions. C-terminal region specific antibody titers have been shown to be associated with protection in RTS,S trial participants, although exposure to *Plasmodium falciparum* strains mismatched to 3D7, significantly reduces protective potency^8-11,21^. The central repeat region and junctional region have been shown to be the target of highly protective monoclonal antibodies in small animal malaria challenge models or in controlled human malaria infection models ^22-27^. Additionally, RTS,S immunogenicity studies showed that the NANP central repeat region is the immunodominant B-cell epitope ^12^. The N-terminal region of CSP (ntCSP) is a disordered region containing Region I (RI), a cystine protease cleavage site ^28^. Subsequently, RI is hypothesized to undergo proteolytic cleavage in response to sporozoite binding to hepatocytes in order to facilitate invasion ^19,28^. Cleavage of this site has been shown to be vital for sporozoite entry into hepatocytes, making it a potential antibody target^28,29^. The N-terminal region has also been identified as a target for complement-fixing mAbs while also sequestering C4b-binding protein, reducing complement fixation and inactivation of sporozoites^30,31^. While mAbs targeting the NANP central repeat region, junctional region, and C-terminal region have been characterized, the protective role of ntCSP mAbs remains understudied.

Historically, mAb5D5 has been the only published and well characterized ntCSP specific monoclonal antibody (mAb)^32,33^. The role of mAb5D5 remains undefined as studies have reported both protective and non-protective in-vivo results^32,33^. Recently, one alternative ntCSP specific mAb, MAD2-6, was discovered and showed protective capabilities, revitalizing the interest in the ntCSP epitope^34^. Still, the lack of an expanded number of diverse and well characterized ntCSP-specific mAbs leaves the protective role of ntCSP-specific immune responses unclear. Here we aimed to characterize mAbs that have specificity for multiple epitopes within the N-terminal region of CSP. C57BL/6 mice underwent an immunization series with a full length ntCSP peptide conjugated to keyhole limpet hemocyanin (KLH) paired with the Sigma Adjuvant System. Utilizing antigen specific B-cell sorting, we selected seven ntCSP mAbs with desirable binding profiles to full length CSP. We then identified their distinct epitope within ntCSP through a peptide walking assay utilizing short overlapping ntCSP peptides.

## Results

### An N-terminal CSP immunogen elicits a CSP specific immune response

To induce an N-terminal CSP-specific immune response in a murine system, we employed a three-dose vaccine regimen (Fig. 1A). Mice were euthanized, and various tissues were isolated. Our immunogen was designed by conjugating amino acid residues 26-100 of *PfCSP* to KLH (Fig. 1B). Mice received intraperitoneal injections at weeks 0, 4, and 12, with intermittent blood draws for determining CSP-specific IgG titers. Recombinant shortened CSP (rsCSP)^35^ was utilized to approximate CSP-specific titers in the serum of immunized animals. CSP specific titers were variable between animals in this study. On average, CSP titers peaked one week after the second boost at 20 μg/mL, stabilizing at approximately 2 μg/mL thereafter (Fig. 1C). For antigen-specific B-cell sorting, splenocytes, PBMCs, and lymphocytes were pooled, and rsCSP was used as a sorting bait. B-cell receptor sequences were obtained through the 10X Genomics VDJ pipeline. All successfully paired IgGs were expressed and screened for N-terminal CSP specificity using a biotinylated N-terminal peptide (Fig. 1B).

**Figure 1:**
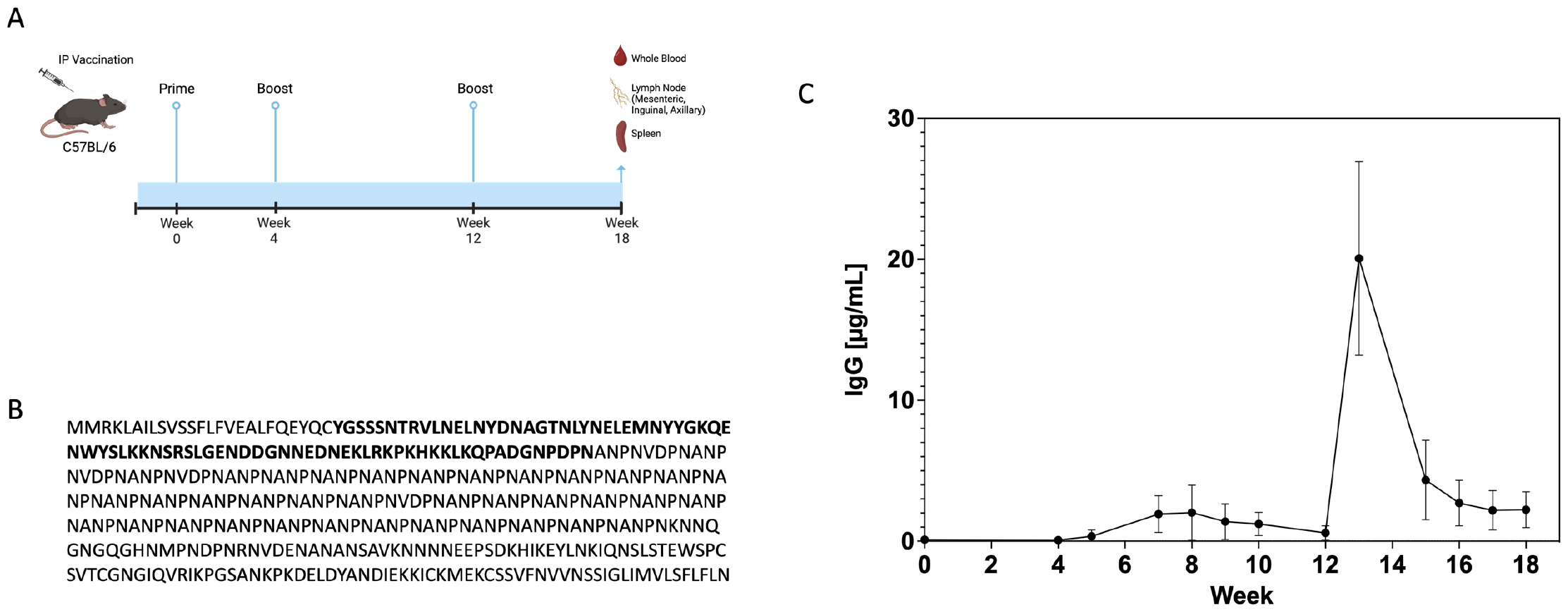
Vaccination with an N-terminal CSP induces antigen specific B-cell responses. A) C57BL/6 mice were immunized with an N-terminal CSP immunogen at weeks 0, 4, and 12. (B) The immunogen was synthesized using *PfCSP* strain 3D7 aa26-100 and conjugated at its N-terminus to KLH. (C) Circulating concentrations of CSP-specific IgG titers were interpolated using a standard curve in a multiplex ELISA format.

### Three B-cell epitope regions are identified in the N-terminus of *Plasmodium falciparum* CSP

To determine the epitope specificities of deselected mAbs, we employed a set of 15 overlapping amino acid peptides spanning residues 26-100 of *PfCSP* in an ELISA format(Fig2A). This peptide walking assay revealed mAb specificities to five different peptides, revealing three B-Cell epitope region bins termed ntCSP-A, ntCSP-B, and ntCSP-C(Fig2A). The murine antibodies mAb5D5^33^ and mNCSP6 target peptides 14 and 16 respectively, placing them in the ntCSP-A bin (Fig2A, Fig2B). This epitope region contains the highly conserved Region I (RI) cysteine protease cleavage site and has been characterized previously. mNCSP9 and mNCSP10 make up the ntCSP-B bin, targeting peptide 8 as their dominant epitope with some specificity to peptide 7 and to a lesser extent peptide 15(Fig2A). The mAbs mNCSP2, mNCSP13, mNCSP14, and mNCSP27 target peptides 1 and 2, making up epitope ntCSP-C(Fig2A, Fig2C). ntCSP-B and ntCSP-C represent two previously undefined B-cell epitope regions located upstream of RI. Affinities of all newly discovered mAbs, along with mAb5D5, were determined for an ntCSP peptide, junctional peptide, repeat motif peptide, and C-terminal CSP peptide using surface plasmon resonance. All mAbs exhibited nanomolar affinity to the N-terminal CSP peptide and showed no specificity for non-N-terminal *PfCSP* epitopes.

**Figure 2:**
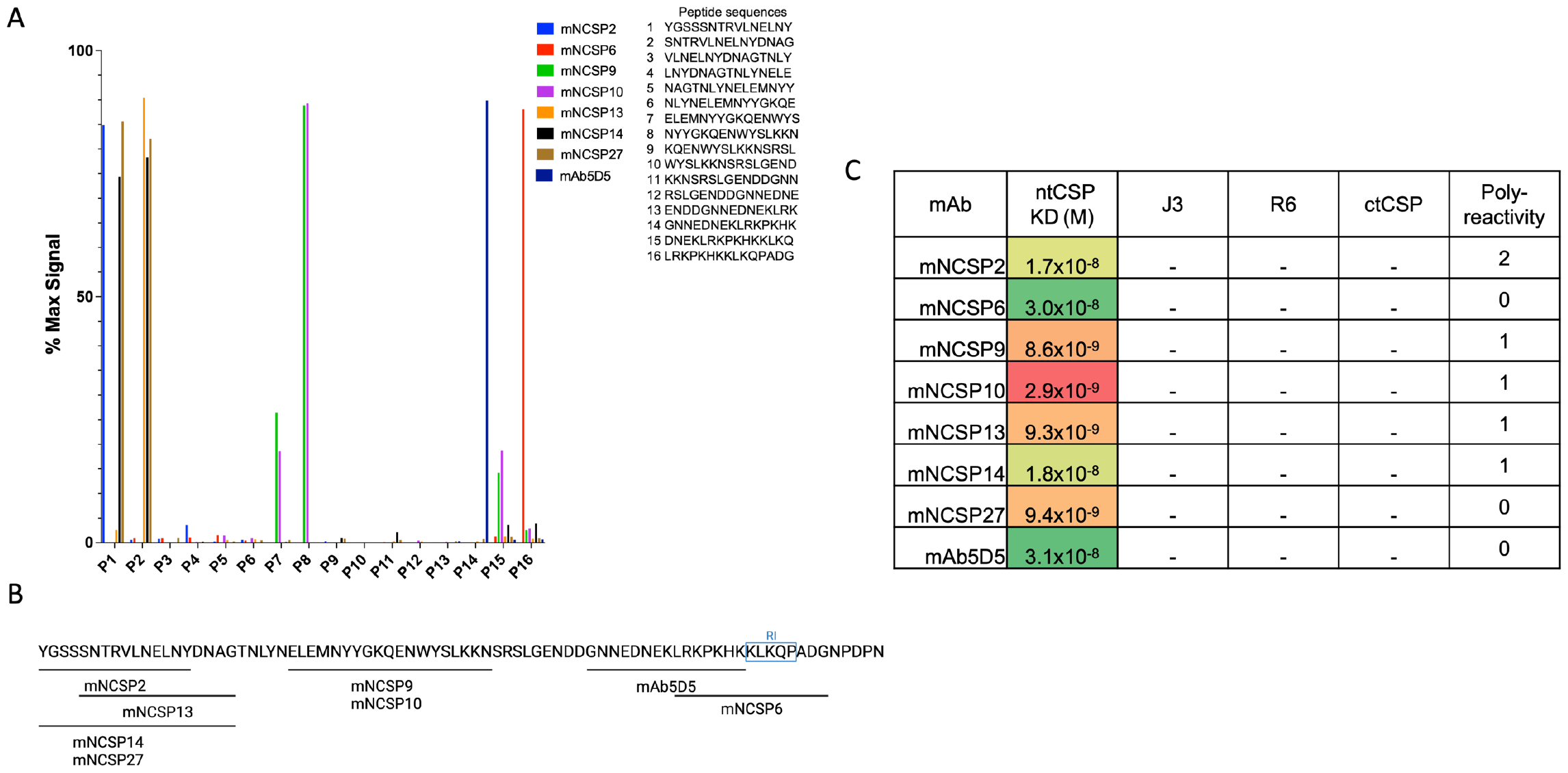
N-terminal CSP specific B-cell responses include three non-overlapping distinct epitope regions. (A) A peptide walking assay was used to determine the approximate epitopes of each mAb. Short 15aa overlapping peptides were used in an ELISA format. (B) Approximate binding locations of each mAb are shown with RI outlined in blue(created with Biorender.com). (C) A Biacore 8K was used to determine the KD affinities to N-terminal CSP (ntCSP), a junctional peptide (J3), repeat motif peptide (R6), and C-terminal CSP peptide (ctCSP). (-) denotes no binding. Polyreactivity was determined using the Hep-2 kit from Hemagen and was scaled from 0-4 where 0 represents negative signal and low polyreactivity and 4 represents high signal and high polyreactivity.

All antibodies underwent screening for polyreactivity to human epithelial type 2 cells through indirect immunofluorescence assays. Antibody polyreactivity was quantified based on the manufacturer’s “criteria for grading fluorescence intensity” and compared to positive and negative controls. mNCSP6, mNCSP27, and mAb5D5 exhibited grade 0 polyreactivity, with fluorescence intensities similar to the negative control. mNCSP9, mNCSP10, mNCSP13, and mNCSP14 received grade 1 polyreactivity, displaying detectable fluorescence signals, although dull and lacking sharpness. Notably, mNCSP2 received the highest polyreactivity grade of 2, demonstrating clearly detectable fluorescence, albeit with lower intensity compared to the positive control.

## Discussion

In this study we characterized eight murine monoclonal antibodies isolated from C57BL/6 mice immunized with an immunogen containing the N-terminal region of *PfCSP*. Murine polyclonal sera targeting the region upstream of RI has been generated in the past to analyze CSP cleavage events^19,28^. Our peptide walking assay indicates the presence of two epitope regions upstream of RI in the N-terminus of CSP in addition to the previously defined RI epitope^33,34^. The ntCSP-A bin is comprised of 2 unique binders, mNCSP6 and mAb5D5, which target peptides containing RI and slightly upstream of RI, respectively. Antibodies targeting this epitope have been shown to have some inhibitory effects in a small animal liver invasion assay^32-34^. However, the true level of protection granted by mAbs in this region has yet to be established. The ntCSP-B bin is made up of two very similar mAbs, mNCSP9 and mNCSP10, targeting the middle of the N-terminus of CSP. The protective qualities of mAbs targeting this region have not yet been elucidated, although it has been shown that this region serves to sequester C4bp, effectively inactivating complement components leading to increased sporozoite fitness^31^. The final bin, ntCSP-C, targeted by mNCSP2, mNCSP13, mNCSP14, and mNCSP27 contains three unique binding sets. The first is comprised of mNCSP2, with binding specificity to peptide 1 only, indicating its epitope is the furthest upstream of RI. The next is comprised of mNCSP13 which exhibits high specificity to peptide 2 alone. The final epitope region is comprised of mNCSP14 and mNCSP27 which spans both peptides 1 and 2.

Overall, our findings reveal antibody binding to three N-terminal CSP epitope regions, two of which are previously undescribed. The vital role of the N-terminal region of CSP plays in sporozoite development and hepatocyte invasion makes it an interesting target for monoclonal antibodies. The protective qualities of the novel mAbs described here have yet to be determined. These insights into the variety of B-cell targets in the N-terminal region promote their further investigation in passive immunization studies and targeted immunogen vaccine studies.

## Methods

### Immunization, B-cell Isolation, and B-cell sequencing

*Plasmodium falciparum* 3D7 CSP was used as a reference to generate an N-terminal immunogen conjugated to KLH (Thermo Fisher Scientific). Murine studies were performed under approved protocol 22-0007. Female C57BL/6 mice 8 weeks of age were immunized through intraperitoneal injection at weeks 0, 4, 12 with 20μg of conjugated protein suspended in the Sigma adjuvant system. Mice were euthanized and tissues were isolated at week 18 for B-cell isolation. A biotinylated rsCSP was conjugated to a streptavidin fluorophore and used as a bait for B-cell sorting. Total antigen specific B cells were collected and run through the 10X Genomics murine VDJ protocols. Samples were sequenced using an Illumina MiSeq system.

### Antibody production

Antibody heavy chain (HC) and light chain (LC) sequence pairs generated from 10X Genomics protocols were synthesized (Geneart, Thermo Fisher Scientific) and cloned into expression vectors containing an IgG_1_ HC or LC constant region. HC and LC constructs were transiently expressed with the Expi293 Expression System (Thermo Fisher). After 5 days, supernatants were harvested and IgG was purified on Protein G Sepharose (GE Healthcare)

### Peptide walking assay

Unconjugated streptavidin (Jackson ImmunoResearch 016-000-084) was coated onto high-binding 384-well plates (Corning 3700) at 2μg/mL overnight at 4°C. Plates were washed and coated with biotinylated 15aa peptides (InnoPep) at 2μg/mL for 1hr at RT. Plates were then washed and blocked with 3% BSA in PBS for 1 hr at RT. After blocking, plates were washed and coated with our mAbs of interest and incubated at RT for 1hr. Plates were then washed and coated with alkaline phosphatase conjugated anti-mouse IgG Fc_y_ secondary (Jackson ImmunoResearch 115-055-008) at 1:2000 dilution and incubated at RT for 1hr. Plates were washed for a final time before developing with alkaline phosphate substrate (Sigma Aldrich S0942). Plates were read at OD_405nm_ and graphed in Prism 10 (version 10.1.1). OD_405nm_ of each mAb was normalized to that mAb’s max signal for binding to rsCSP.

### Surface Plasmon Resonance

Affinity experiments were performed on a Biacore 8k at 25°C at a flow rate of 30μL/min in a mobile phase of HBS-EP+ [0.01 M HEPES (pH 7.4), 0.15 M NaCl, 3 mM EDTA, 0.0005% (v/v) Surfactant P20] for all experiments. mAb was injected over a protein G series S chip (Cytiva) followed by a wait period for normalization of response units (RU). A concentration series of an N-terminal CSP, junctional CSP, repeat motif CSP, or C-terminal CSP peptides were injected across the antibody and control surface for 120 seconds, followed by a 500 second dissociation phase. An injection with 10 mM Glycine-HCl pH 1.5 was used for chip regeneration. Kinetic analysis of each reference subtracted injection series was performed using BIAEvaluation software (Cytiva).

### Polyreactivity

Reactivity to human epithelial type 2 (HEp2) cells was determined by indirect immunofluorescence on HEp2 slides (Hemagen 902360) according to manufacturer’s instructions. Monoclonal antibody was diluted at 50 μg/mL in PBS and then incubated onto immobilized HEp2 slides for 30 min at room temperature. After washing 2X with PBS, one drop of fluorescein isothiocyanate (FITC)-conjugated goat anti-mouse IgG was added onto each well and incubated in the dark for 30 min at room temperature. After washing, the coverslip was added to HEp2 slide with glycerol and the slide was photographed on a Nikon fluorescence microscope to measure FITC signal. Cross-reactivity profiles were determined through the presence of FITC signals.

## References

1 WHO. World Malaria Report 2023. (2023).

2 Lubell, Y. et al. Artemisinin resistance--modelling the potential human and economic costs. Malar J 13, 452 (2014).

3 World Health Organization. Artemisinin resistance and artemisinin-based combination therapy efficacy: status report. (World Health Organization, 2018).

4 World Health Organization. Global Technical Strategy for Malaria 2016–2030. (World Health Organization, 2015).

5 Rts, S. C. T. P. First results of phase 3 trial of RTS,S/AS01 malaria vaccine in African children. The New England journal of medicine 365, 1863–1875 (2011).

6 Olotu, A. et al. Seven-year efficacy of RTS, S/AS01 malaria vaccine among young African children. The New England journal of medicine 374, 2519–2529 (2016).

7 Casares, S., Brumeanu, T. D. & Richie, T. L. The RTS,S malaria vaccine. Vaccine 28, 4880–4894 (2010). 10.1016/j.vaccine.2010.05.033

8 Dobano, C. et al. Concentration and avidity of antibodies to different circumsporozoite epitopes correlate with RTS,S/AS01E malaria vaccine efficacy. Nature communications 10, 2174 (2019).

9 Chaudhury, S. et al. Breadth of humoral immune responses to the C-terminus of the circumsporozoite protein is associated with protective efficacy induced by the RTS,S malaria vaccine. Vaccine 39, 968–975 (2021).

10 Suscovich, T. J. et al. Mapping functional humoral correlates of protection against malaria challenge following RTS,S/AS01 vaccination. Sci Transl Med 12 (2020). 10.1126/scitranslmed.abb4757

11 S Moses Dennison, M. R., Milite Abraha, Rachel L Spreng, Ulrike Wille-Reece, Sheetij Dutta, Erik Jongert, S Munir Alam, Georgia D Tomaras. Magnitude, Specificity, and Avidity of Sporozoite-Specific Antibodies Associate With Protection Status and Distinguish Among RTS,S/AS01 Dose Regimens. Open Forum Infectious Disease 8 (2021). 10.1093/ofid/ofaa644

12 White, M. T. et al. Immunogenicity of the RTS,S/AS01 malaria vaccine and implications for duration of vaccine efficacy: secondary analysis of data from a phase 3 randomised controlled trial. The Lancet infectious disease 15, 1450–1458 (2015).

13 Laurens, M. B. RTS,S/AS01 vaccine (Mosquirix): an overview. Hum Vaccin Immunother 16, 480–489 (2020). 10.1080/21645515.2019.1669415

14 Datoo, M. S. et al. Efficacy and immunogenicity of R21/Matrix-M vaccine against clinical malaria after 2 years’ follow-up in children in Burkina Faso: a phase 1/2b randomised controlled trial. The Lancet. Infectious diseases 22, 1728–1736 (2022). 10.1016/S1473-3099(22)00442-X

15 Datoo, M. S. et al. Efficacy of a low-dose candidate malaria vaccine, R21 in adjuvant Matrix-M, with seasonal administration to children in Burkina Faso: a randomised controlled trial. Lancet 397, 1809–1818 (2021). 10.1016/S0140-6736(21)00943-0

16 Zavala, F. et al. Rationale for development of a synthetic vaccine against Plasmodium falciparum malaria. Science 228, 1436–1440 (1985).

17 Doud, M. B. et al. Unexpected fold in the circumsporozoite protein target of malaria vaccines. Proceedings of the National Academy of Sciences of the United States of America 109, 7817–7822 (2012).

18 Pradel, G., Garapaty, S. & Frevert, U. Proteoglycans mediate malaria sporozoite targeting to the liver. Mol Microbiol 45, 637–651 (2002). 10.1046/j.1365-2958.2002.03057.x

19 Coppi, A. et al. Heparan sulfate proteoglycans provide a signal to Plasmodium sporozoites to stop migrating and productively invade host cells. Cell host & microbe 2, 316–327 (2007). 10.1016/j.chom.2007.10.002

20 Beutler, N. et al. A novel CSP C-terminal epitope targeted by an antibody with protective activity against Plasmodium falciparum. PLoS Pathog 18, e1010409 (2022). 10.1371/journal.ppat.1010409

21 Neafsey, D. E. et al. Genetic Diversity and Protective Efficacy of the RTS,S/AS01 Malaria Vaccine. The New England journal of medicine 373, 2025–2037 (2015). 10.1056/NEJMoa1505819

22 Kisalu, N. K. et al. A human monoclonal antibody prevents malaria infection by targeting a new site of vulnerability on the parasite. Nature medicine 24, 408 (2018).

23 Tan, J. et al. A public antibody lineage that potently inhibits malaria infection through dual binding to the circumsporozoite protein. Nature medicine 24, 401 (2018).

24 Imkeller, K. et al. Antihomotypic affinity maturation improves human B cell responses against a repetitive epitope. Science 360, 1358–1362 (2018).

25 Oyen, D. et al. Structural basis for antibody recognition of the NANP repeats in Plasmodium falciparum circumsporozoite protein. Proceedings of the National Academy of Sciences of the United States of America 114, E10438–E10445 (2017).

26 Pholcharee, T. et al. Structural and biophysical correlation of anti-NANP antibodies with in vivo protection against P. falciparum. Nat Commun 12, 1063 (2021).

27 Lyke, K. E. et al. Low-dose intravenous and subcutaneous CIS43LS monoclonal antibody for protection against malaria (VRC 612 Part C): a phase 1, adaptive trial. The Lancet. Infectious diseases 23, 578–588 (2023). 10.1016/S1473-3099(22)00793-9

28 Coppi, A., Pinzon-Ortiz, C., Hutter, C. & Sinnis, P. The Plasmodium circumsporozoite protein is proteolytically processed during cell invasion. The Journal of experimental medicine 201, 27–33 (2005). 10.1084/jem.20040989

29 Coppi, A. et al. The malaria circumsporozoite protein has two functional domains, each with distinct roles as sporozoites journey from mosquito to mammalian host. The Journal of experimental medicine 208, 341–356 (2011). 10.1084/jem.20101488

30 Kurtovic, L., Drew, D. R., Dent, A. E., Kazura, J. W. & Beeson, J. G. Antibody Targets and Properties for Complement-Fixation Against the Circumsporozoite Protein in Malaria Immunity. Frontiers in immunology 12, 775659 (2021). 10.3389/fimmu.2021.775659

31 Khattab, A. et al. Hijacking the human complement inhibitor C4b-binding protein by the sporozoite stage of the Plasmodium falciparum parasite. Frontiers in immunology 13, 1051161 (2022). 10.3389/fimmu.2022.1051161

32 Thai, E. et al. A high-affinity antibody against the CSP N-terminal domain lacks Plasmodium falciparum inhibitory activity. The Journal of experimental medicine 217 (2020). 10.1084/jem.20200061

33 Espinosa, D. A. et al. Proteolytic cleavage of the Plasmodium falciparum circumsporozoite protein is a target of protective antibodies. J Infect Dis 212, 1111–1119 (2015).

34 Tan, J. et al. Functional human IgA targets a conserved site on malaria sporozoites. Sci Transl Med 13 (2021). 10.1126/scitranslmed.abg2344

35 Schwenk, R. et al. IgG2 antibodies against a clinical grade Plasmodium falciparum CSP vaccine antigen associate with protection against transgenic sporozoite challenge in mice. PloS one 9, e111020 (2014). 10.1371/journal.pone.0111020

